# A new nitrogen biochemical process: Oxidization of ammonium into nitrogen gas using nitrate

**DOI:** 10.1101/2025.06.12.659240

**Authors:** Xintao Lv, Shujun Zhang, Pengchao Gu, Xiaoyu Han, Quan Li, Jing Huang, Zhiming Qu, Dan Zhao, Zhibin Wang, Kun Li, Cong Wang

## Abstract

The nitrifying bacteria oxidize ammonium into nitrite or nitrate under aerobic conditions, while the anaerobic ammonia-oxidizing bacteria utilize nitrite to oxidize ammonium into nitrogen gas under anaerobic conditions. There is no biochemical process that can directly oxidize ammonium into nitrogen gas using nitrate under anaerobic conditions. In this study, mature anaerobic ammonium oxidation (anammox) granular sludge was inoculated in an anaerobic nitrogen-removal system, while nitrate and ammonium were used as the influent substrates. The experiment was conducted for 537 days at a temperature of 30-32°C and a pH of 8.0-9.0. The transformation from nitrite-anammox to nitrate-anammox was stably achieved on the 350^th^ day. The experimental results showed that, nitrate directly oxidized ammonium into nitrogen gas under anaerobic conditions. The produced gas consisted of 96.3% nitrogen and 3.7% carbon dioxide. The removal ratio of ammonium to nitrate was approximately 1.67:1, and the total inorganic nitrogen removal rate reached up to 290.20 mg/(L·d). Unlike the nitrite-anammox reaction, this reaction was accompanied by an acid production process, which caused a decrease in pH. When the substrate was changed to nitrite and ammonium, the total nitrogen removal rate of only 1.1-1.3 mg/(L·d) was achieved. This new biochemical reaction of nitrogen was defined as nitrate-anammox. The study reveals a new pathway of nitrogen transformation, providing novel insights into the global nitrogen cycle.

## 1 Introduction

Ammonia oxidation is an important component of nitrogen cycle, which is widely prevalent in nature, as well as in industrial and agricultural environments (Hutchins & Capone, 2022; Kuypers et al., 2018; Lehnert et al., 2021). At the end of the 19^th^ century, scientists discovered nitrifying bacteria. For approximately 100 years thereafter, scholars believed that nitrifying bacteria were the only microorganisms capable of oxidizing 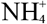 into 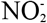 and 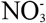, and the molecular oxygen must be involved in this process. However, this belief was completely overturned after the discovery of anaerobic ammonium oxidation (anammox) reaction, which uses 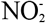 as the electron acceptor and involves functional microorganisms (Kuypers et al., 2018; Mulder et al., 1995). Anaerobic ammonium oxidizing bacteria (AnAOB) are widely distributed around the world and are an important component of nitrogen cycle. Anammox may contribute 30%-50% of N_2_ in the ocean (Devol, 2003; Kartal et al., 2010). The anammox reaction can be represented as follows:

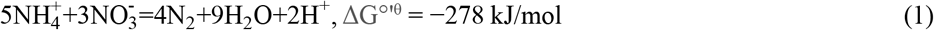

This equation indicates that the reaction between 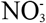 and 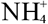 can proceed spontaneously.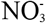 is more common in nature than 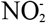 (Stüeken et al., 2024), and it is more likely to coexist with 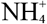. However, so far, there is no known biochemical process that oxidizes ammonia nitrogen into nitrogen gas using nitrate.

In this study, anammox granular sludge was collected from a wastewater treatment reactor and used as the inoculum. 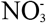 and 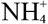 were used as the influent substrates and supplied as the components of test water prepared manually. The treatment experiment was conducted for 537 days at a temperature range of 30-32°C and a pH range of 8.0-9.0, and the functional microorganisms capable of 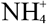 oxidization using 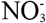 were successfully enriched. The study reveals a novel biochemical process which oxidizes ammonium into nitrogen gas using nitrate.

## 2 Materials and methods

### 2.1 Experimental devices, test water preparation, and characteristics of inoculum sludge

The experiments were conducted using a sequencing batch reactor (SBR) (Fig. 1) and an up-flow anaerobic sludge blanket (UASB) reactor (Fig. 2), with effective volumes of 15 L and 7 L, respectively. Both reactors were made of organic glass.

**Figure 1.**
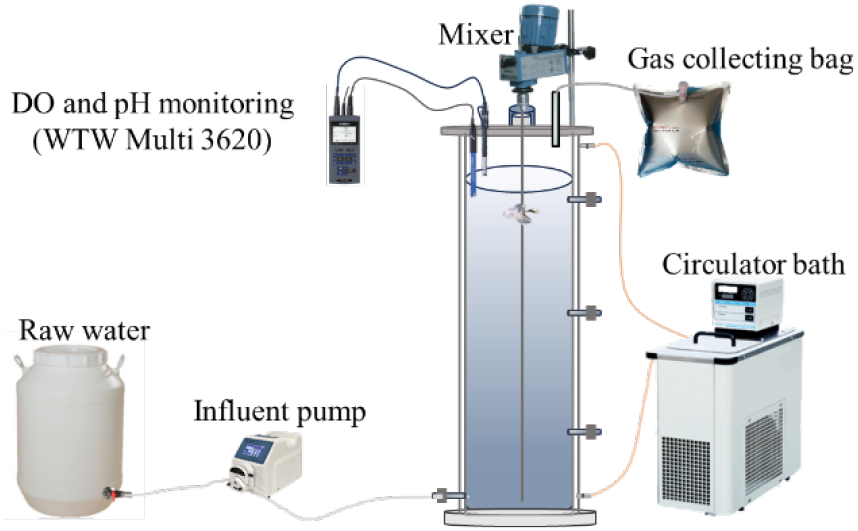
SBR.

**Figure 2.**
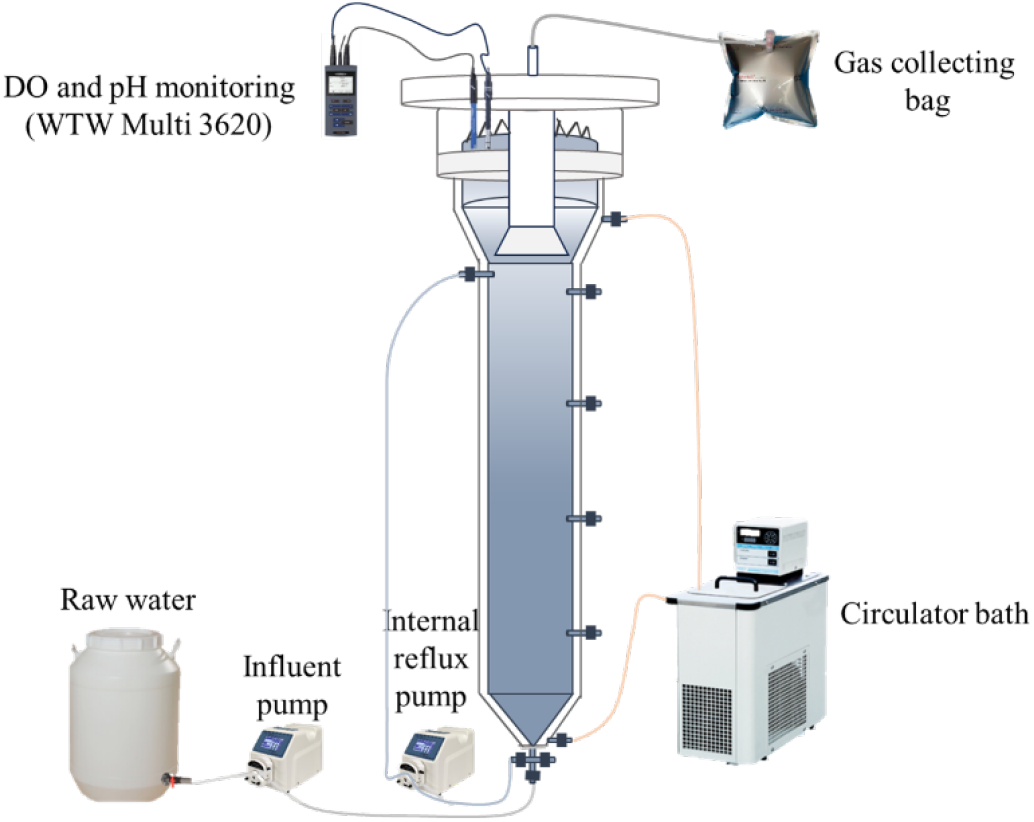
UASB reactor.

The test water was prepared manually. The concentrations of 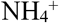 and 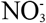 were adjusted by adding NH_4_HCO_3_ solution (100 g/L) and NaNO_3_ (100 g/L), respectively. The pH was adjusted by 0.1 mol/L HCl and NaHCO_3_ (0.4 g/L). In addition, minerals MgSO_4_·7H_2_O (0.3 g/L), CaCl_2_·2H_2_O (0.18 g/L), KH_2_PO_4_(0.0272 g/L) were added. Trace element I (1 mL) and trace element II (0.5 mL) solutions were added per liter of prepared water. The compositions of both trace element solutions are shown in Table 1. The anammox granular sludge was taken from the wastewater treatment plant in Gaobeidian, Beijing, and used as the inoculum. The sludge had a particle size of 500-1500 μm and a good nitrogen removal function. The nitrogen removal load was 700-1000 mg/(L·d). After inoculation, the sludge concentration in the incubator was 13,000 mg/L.

**Table 1.**
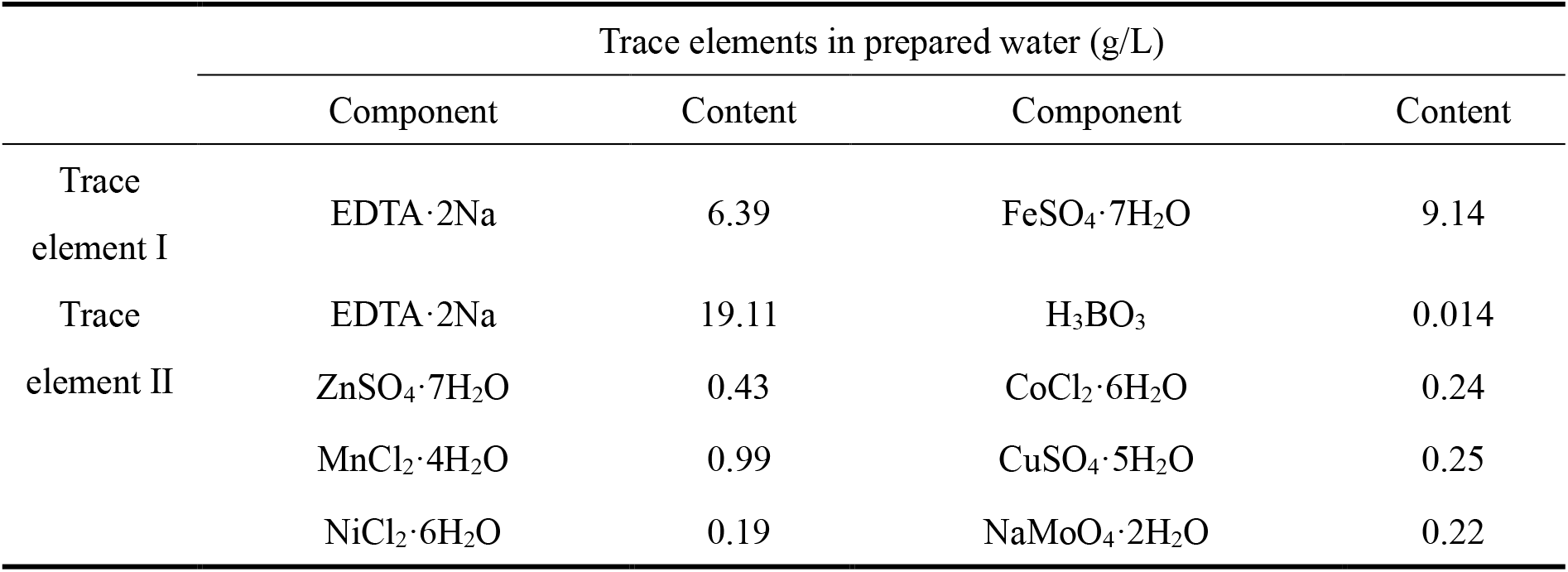
Compositions of trace element solutions.

### 2.2 Experiment control conditions

SBR control conditions: The temperature in the reactor was controlled at 30-32°C through a water bath. Dissolved oxygen (DO) was not detected. At regular intervals of 7-14 d, 67% of the water in SBR was replaced with fresh test water. The influent pH was 8.0-9.0, and the mixing speed was 100 rpm.

UASB control conditions: At 446 d, the sludge from the SBR was transferred to the UASB reactor. The temperature in the UASB reactor was controlled at 30-32°C using a water bath. The influent pH was 8.0-9.0, while the DO concentration was not detected. The hydraulic retention time (HRT) was 0.5-3 days.

The experiment was divided into three stages. In Stage I (0-223 d), SBR operation was conducted. The concentrations of 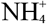 and 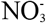 in the influent were 47.3-70.6 mg/L and 40-50.3 mg/L, respectively. Stage II (224-445 d) also included SBR operation, with influent 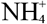 and 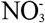 concentrations of 57.3-103.6 mg/L and 44.5-63.2 mg/L, respectively. In Stage III (446-537 d), the treatment mode was changed to continuous flow, by transferring the sludge from SBR to UASB reactor. The concentrations of 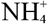 and 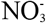 in the influent were maintained at 48.7-125.0 mg/L and 36.8-70.2 mg/L, respectively.

### 2.3 Calculations

HRT was determined by using the following equation:

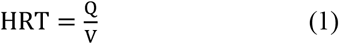

Where Q denotes the influent flow rate (L/h) and V denotes the effective volume of reactor (L). The total inorganic nitrogen concentration (C_TIN_) was calculated as follows:

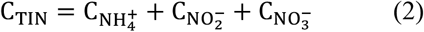

Where C_NH+ 4_ denotes the 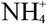 concentration; C_NO-2_ denotes the 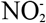 concentration; and C_NO-3_ denotes the 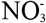 concentration. All concentrations were presented as mg/L.

The total inorganic nitrogen removal rate (R_TIN_) was calculated using the following equation.

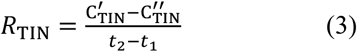

Where 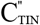 denotes the total inorganic nitrogen concentration at the end of reaction (mg/L), while *t*_1_ and *t*_2_ denote the beginning time and the end time of reaction, respectively.

The ratio of 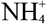 concentration removal to 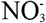 concentration removal 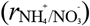 was determined as follows:

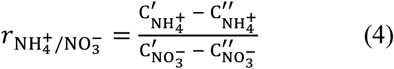

Where 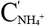 and 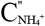 denote the initial and final 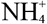 concentrations, respectively, while 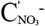 and denote the initial and final 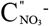 concentrations, respectively. All concentrations were measured as mg/L.

### 2.4 Analytical methods

The samples collected from the reactors were filtered through a 0.45-μm filter. The concentrations of 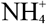, 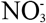, and 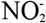 in the sludge were determined using the standard methods (Eaton et al., 1966). The temperature of sludge samples was measured using a portable WTW-Multi 3420 meter (WTW, Germany). Gas detection was performed using a portable gas analyzer (Biogas ES 5K, United Kingdom).

## 3 Results and discussion

### 3.1 Changes in the water quality indicators during the experiment

Fig. 3 shows the changes in the water quality indicators, including 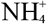, 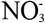, 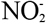, pH, *R*_TIN_, and 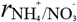 during the experiment. To verify whether the reaction occurring in the reactor is a biochemical reaction dominated by nitrate-anammox, the concentrations of 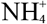 and 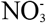 in the influent were changed during 413-419 d, 430-436 d, 437-443 d, and 510-516 d, respectively. In Stage I (0-223 d), the concentrations of 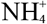 and 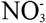 in the influent were 47.3-70.6 mg/L and 40.0-50.3 mg/L, respectively, while 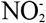 was not detected. As the reactor operation continued, the *R*_TIN_ decreased from 6.5 mg/(L·d) to 0.3 mg/(L·d). Correspondingly, the concentration of 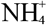 in the effluent increased from 17.6 mg/L to 67.6 mg/L, while 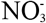 concentration increased from 0.2 mg/L to 47.5 mg/L. In the initial stage of treatment, the pH of sludge increased, while *r*_NH+ 4/NO-3_ initially increased from 0.68 to 2.48 and then decreased to 1.04.

**Figure 3.**
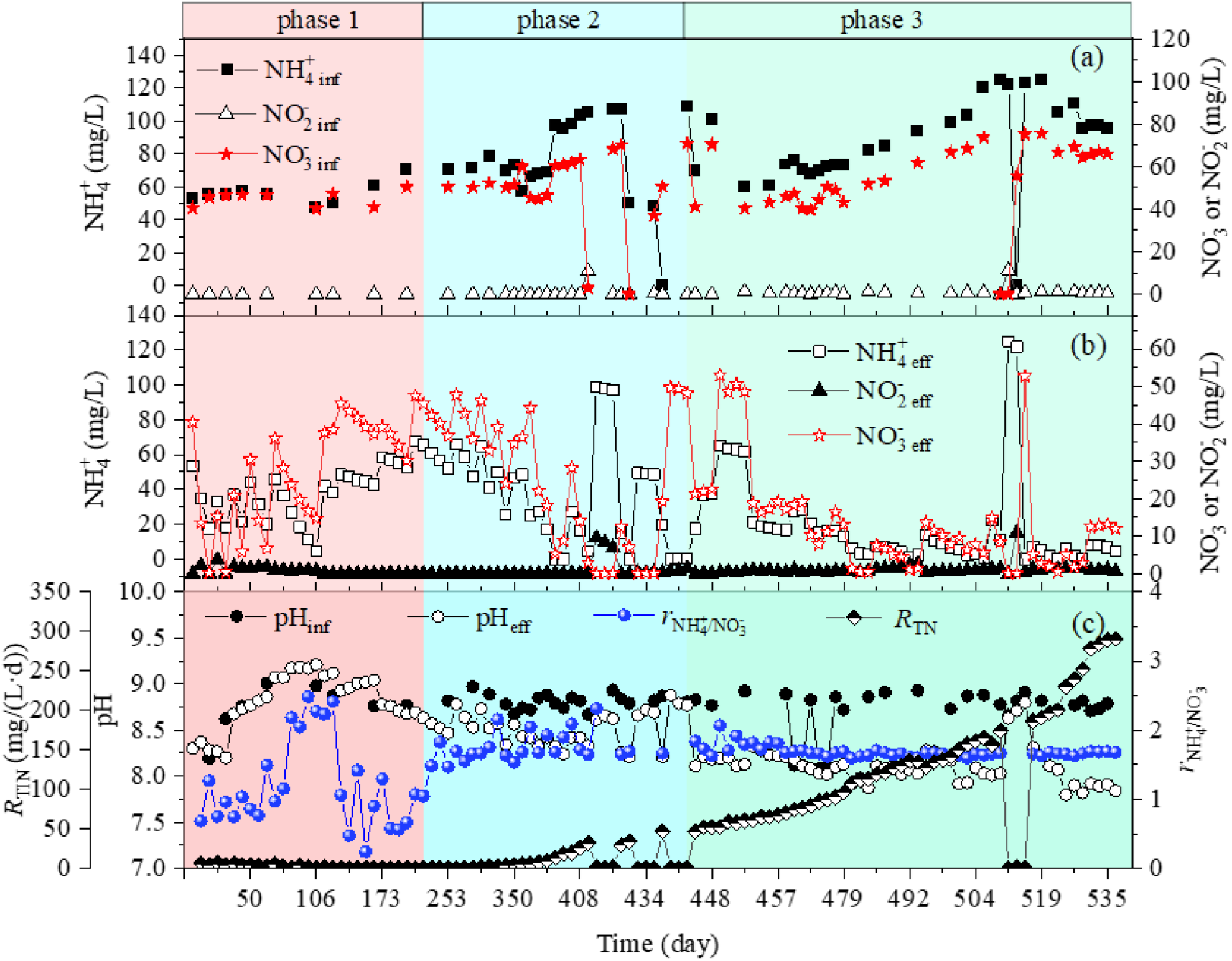
Changes in 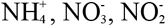, pH, *R*_TIN_, and 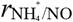 during the treatment process.

In Stage I, denitrifying bacteria in the reactor used the organic matter released after the decay of anammox granular sludge to reduce a portion of 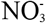 to 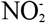 (short-term nitrogen removal).

Subsequently, the AnAOB used this generated 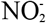 to oxidize 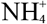 into N_2_ (nitrite-anammox reaction), which led to a decrease in C_TIN_ and an increase in pH. With the continuous consumption of organic matter in the reactor, the short-term nitrogen removal and nitrite-anammox reactions gradually weakened, leading to a significant decrease in *R*_TIN_.

During Stage II (224-445 d), the concentrations of 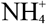 and 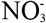 in the influent were 57.3-103.6 mg/L and 44.5-63.2 mg/L, respectively, while 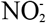 was not detected in the influent.

The concentrations of 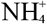 and 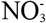 in the effluent gradually decreased from 67.6 mg/L to 4.2 mg/L and from 47.5 mg/L to 2.9 mg/L, respectively. The *R*_TN_ gradually increased from 0.43 mg/(L·d) to 46.90 mg/(L·d), with an average 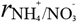 of 1.73. The pH of sludge in the reactor showed a continuous decreasing trend, and the decrease range gradually expanded from 0.01 to 0.1.

In Stage III (446-537 d), the treatment mode was changed from SBR to UASB and the HRT was gradually decreased from 7-14 d to 0.5-3 d. The concentrations of 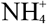 and 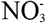 in the influent water were 48.7-125.0 mg/L and 36.8-70.2 mg/L, respectively. The ratio of 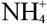 to 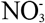 in the influent was maintained at 1.5-1.7. As the water inflow gradually increased, *R*_TIN_ also increased from 31.94 mg/(L·d) to 290.20 mg/(L·d). The pH of sludge continued to decrease, and the decrease range gradually increased from 0.10 to 0.95. The 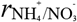 value remained stable in the range of 1.6-1.8.

The long-term stable experimental results of Stage II and Stage III were very consistent with the reaction shown in Eq. (1), with 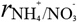 being close to the stoichiometric ratio of 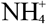 to 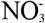 (5/3≈1.67). At the same time, the continuous decrease in pH was also consistent with the acid production shown in reaction (1) and different from the pH increase characteristics shown in nitrite-anammox reaction (2) below:

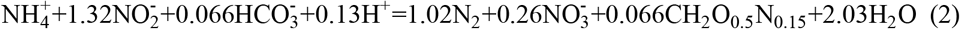

It can be concluded that the biochemical reaction in the reactor was a nitrate-anammox reaction discovered for the first time.

### 3.2 Verification of nitrate-anammox as the dominant biochemical reaction in reactor

To verify the occurrence of nitrate-anammox in the reactor, the influent substrate was adjusted on 415-447 d and 510-519 d, respectively. Fig. 4 shows the variations in the nitrogen removal efficiency of treatment system after adjustments in the influent substrate. On the 415^th^ d, after the influent substrate was changed from 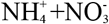 to 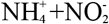, the concentrations of 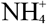 and 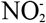 in the effluent decreased from 99.0 mg/L to 96.6 mg/L and from 9.6 mg/L to 9.2 mg/ L, respectively. The pH values of influent and effluent were basically the same. These results indicate that the anammox and nitrogen removal in the reactor greatly weakened compared to the initial stage of treatment. On the 419^th^ d, after changing the substrate from 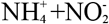 to 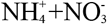, the concentrations of both 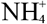 and 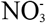 in the effluent decreased significantly. Simultaneously, the pH decreased from 8.84-8.93 to 8.21-8.26. These findings indicate the rapid recovery of nitrate-anammox in the reactor. On the 430^th^ d, the substrate was changed from 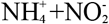to 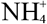. After this modification, the concentration of 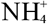 in the effluent decreased from 50.3 mg/L to 48.7 mg/L, while the concentration of 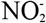 increased from 0 mg/L to 0.4 mg/L. 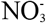 was not detected. This indicated the occurrence of a weak nitrification reaction in the incubator. Furthermore, the nitrate-anammox reaction removed the possible trace amounts of 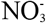 and 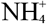 from the influent. On the 436^th^ d, after changing the influent substrate from 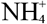 to 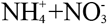, the concentrations of 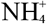 and 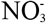 decreased from 48.7 mg/L to 19.4 mg/L and from 36.8 mg/L to 19.2 mg/L, respectively, while the pH decreased from 8.82 to 8.21. These findings indicate the rapid recovery of nitrate-anammox in the reactor. On the 437^th^ d, the substrate was transformed from 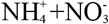 to 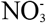. After 6 days of treatment, the concentration of 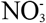 decreased from 50.7 mg/L to 48.1 mg/L, while 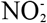 concentration increased from 0 mg/L to 1.4 mg/ L. 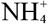 concentration and pH did not show any significant change. These results indicate that only weak short-term nitrogen removal occurred in the reactor during this time, and the nitrogen removal pathways dominated by heterotrophic denitrifying bacteria and AnAOB could be disregarded. On the 443^th^ d, after transformation of substrate from 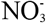 to 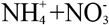, the concentrations of 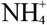 and 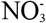, as well as the pH decreased significantly. The nitrogen-anammox functional flora dominated the nitrogen removal process in the reactor during this stage.

**Figure 4.**
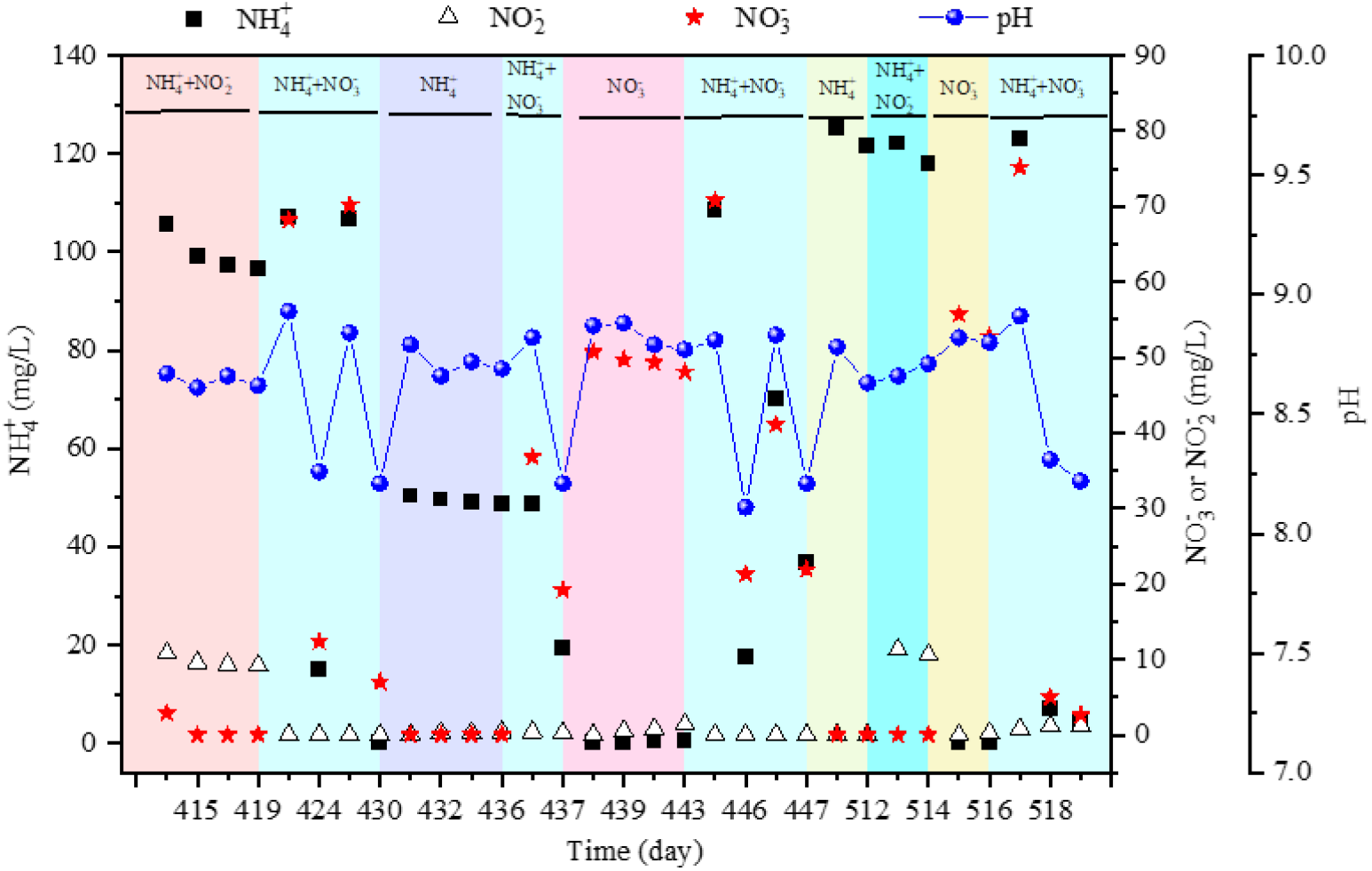
Influence of adjusting the influent substrate on nitrogen removal efficiency during the experiment.

When *R*_TIN_ continued to increase, the influent substrate was successively changed from 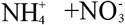 to 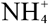, 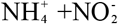, and 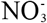 to verify the occurrence of nitrate-anammox again.

Furthermore, the changes in the TIN removal efficiency of system were investigated to determine the main nitrogen removal pathway. On the 510^th^ d, 512^th^ d, and 514^th^ d, the influent substrate was changed from 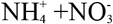 to 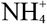, from 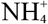 to 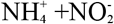, and from 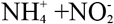 to 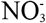, respectively. Under the above three substrate conditions, compared with the influent, the concentrations of 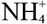, 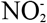, and 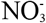 in the effluent were basically similar in comparison to those in the influent, and the pH also did not declined significantly. However, when the substrate was changed from 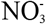 to 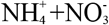 on the 516^th^ d, the concentrations of 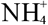 and 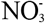 in the effluent, as well as pH decreased significantly. The above experiments once again proved the enrichment of nitrate-anammox functional microbes in the reactor and the dominance of novel nitrogen removal reaction (i.e., 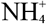 oxidation to nitrogen gas by 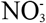) in the system.

## 4 Conclusions and Prospects

The ammonium oxidation process mainly includes aerobic ammonium oxidation and anammox (i.e., nitrite anammox and nitrate anammox). After 537 d of treatment, the nitrate-anammox functional microbes were successfully enriched, and the simultaneous removal of 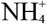 and 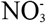 from test water was achieved for the first time. The *R*_TIN_ was as high as 290.2 mg/(L·d), with a 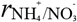 ratio of approximately 1.67. The reaction was an acid-producing process, which led to a decline of 0.95 in the pH of effluent. The discovery of nitrate-anammox will provide novel insights into the global nitrogen cycle pathways. The findings of this study will be helpful in re-evaluating the global nitrogen flux quantitatively. This study will encourage the scientific community to redefine the nitrogen migration/transformation mechanisms in different ecosystems and provide a brand-new path for biological nitrogen removal from wastewater.

## References

Devol, A.H. 2003. Nitrogen cycle: Solution to a marine mystery. Nature, 422(6932), 575–6.

Eaton, A.D., Clesceri, L.S., Greenberg, A.E., Franson, M.A.H. 1966. Standard methods for the examination of water and wastewater. Am J Public Health Nations Health, 56(3), 387–388.

Hutchins, D.A., Capone, D.G. 2022. The marine nitrogen cycle: new developments and global change. Nat Rev Microbiol, 20(7), 401–414.

Kartal, B., Kuenen, J.G., Van Loosdrecht, M.C.M. 2010. Sewage Treatment with Anammox. Science, 328(5979), 702–703.

Kuypers, M.M.M., Marchant, H.K., Kartal, B. 2018. The microbial nitrogen-cycling network. Nat Rev Microbiol, 16(5), 263–276.

Lehnert, N., Musselman, B.W., Seefeldt, L.C. 2021. Grand challenges in the nitrogen cycle. Chem Soc Rev, 50(6), 3640–3646.

Mulder, A., Graaf, A.A.V.D., Robertson, L.A., Kuenen, J.G. 1995. Anaerobic ammonium oxidation discovered in a denitrifying fluidized bed reactor. FEMS Microbiology Ecology, 16(3), 177–183.

Stüeken, E.E., Pellerin, A., Thomazo, C., Johnson, B.W., Duncanson, S., Schoepfer, S.D. 2024. Marine biogeochemical nitrogen cycling through Earth’s history. Nature reviews Earth & environment, 5(10), 732–747.

